# Cognitive Vergence and Pupil Response During Oddball Task are Associated With Alzheimer’s Disease Cerebrospinal Fluid Neurodegenerative Biomarkers

**DOI:** 10.64898/2026.04.10.717637

**Authors:** Ricardo Martínez-Flores, Isabel Martín-Sobrino, Neus Falgàs, Oriol Grau-Rivera, Marc Suárez-Calvet, Carlos Cristi-Montero, Agustín Ibañez, Hans Supèr

**Affiliations:** Institute of Neurosciences, University of Barcelona (UBNeuro), 08035 Barcelona, Spain; Vision and Control of Action Group, Department of Cognition, Development and Educational Psychology, University of Barcelona, 08035 Barcelona, Spain; Alzheimer’s Disease and Other Cognitive Disorders Unit, Neurology Service, Hospital Clínic de Barcelona, Fundació de Recerca Clínic Barcelona-IDIBAPS, Barcelona, Catalonia, Spain; Universitat de Barcelona, Barcelona, Spain; Centro de Investigación Biomédica en Red en Enfermedades Neurodegenerativas (CIBERNED), Madrid, Spain; Barcelonaβeta Brain Research Center (BBRC), Pasqual Maragall Foundation, Barcelona, Spain; Hospital del Mar Research Institute, Barcelona, Spain; Centro de Investigación Biomédica en Red de Fragilidad y Envejecimiento Saludable (CIBERFES), Instituto de Salud Carlos III, Madrid, Spain; Servei de Neurologia, Hospital del Mar, Barcelona, Spain; IRyS Group, Physical Education School, Pontificia Universidad Católica de Valparaíso, Valparaíso, Chile; Center for Interdisciplinary Research in Biomedicine, Biotechnology and Well-Being (CID3B), Pontificia Universidad Católica de Valparaíso, Valparaíso, Chile; Latin American Brain Health Institute (BrainLat), Universidad Adolfo Ibáñez, Santiago, Chile; Global Brain Health Institute, Trinity College Dublin, Dublin, Ireland; Department of Biophysics, School of Medicine, Istanbul Medipol University, 34815 Istanbul, Türkiye; Barcelonaβeta Brain Research Center (BBRC), Pasqual Maragall Foundation, 08005, Barcelona, Spain; Cognitive Neuroscience Center (CNC), Universidad de San Andrés, Buenos Aires, Argentina; Braingaze SL, Mataró, Spain; Catalan Institution for Research and Advanced Studies (ICREA), Barcelona, Spain

**Keywords:** Alzheimer’s disease, CSF biomarkers, Eye-tracking, Locus coeruleus, Attention networks

## Abstract

**Background:** Alzheimer’s disease (AD) can be diagnosed using cerebrospinal fluid (CSF) biomarkers reflecting amyloid and tau pathology. However, it provides no information about functional network status. We aimed to determine whether CSF biomarkers (Aβ42, p-Tau, t-Tau, and Aβ42/p-Tau ratio) are associated with altered stimulus differentiation in vergence and pupil responses during an oddball task, and to evaluate oculomotor metrics as predictors of CSF core AD biomarkers in patients at mild cognitive impairment (MCI) stage.

**Methods:** Thirty-eight participants with abnormal CSF core AD biomarkers at MCI stage completed a visual oddball task while oculomotor responses were recorded. Linear mixed-effects models examined condition × biomarker interactions, controlling for sex, age, and MMSE. Temporal and magnitude features were tested as predictors using linear regression.

**Results:** Higher p-Tau levels were negatively associated with target-distractor differentiation in cognitive vergence (β = -0.035, p < 0.001) and pupil responses (β = - 0.060, p < 0.001). Higher Aβ42 and Aβ42/p-Tau showed positive associations with vergence differentiation but opposite effects on pupil responses. Oculomotor features predicted p-Tau levels (R^2^ = 0.20–0.21).

**Conclusion:** Oculomotor differentiation metrics capture functional signatures of tau-related network dysfunction, positioning them as accessible biomarkers complementing CSF measures for detecting network disruption at MCI stage.

## 1. Introduction

Alzheimer’s disease (AD) represents a progressive neurodegenerative condition characterized by accumulation of specific misfolded proteins (amyloid-beta and tau) can now be detected and staged using biological markers during life ^1^. The worldwide prevalence of AD is rising dramatically, affecting approximately 50 million individuals currently, with epidemiological models projecting this figure could reach 115 million by mid-century ^2^. As populations age globally, dementia has emerged as a major contributor to disability and loss of independence in older adults, emphasizing the urgent need for accessible tools enabling early identification and longitudinal monitoring of disease progression ^3^.

The pathological hallmarks of AD include extracellular amyloid-beta (Aβ) plaques and intracellular neurofibrillary tangles composed of hyperphosphorylated tau protein, detectable through cerebrospinal fluid (CSF) biomarkers: decreased Aβ42 levels reflect amyloid deposition and are among the earliest detectable molecular changes, elevated phosphorylated tau (p-Tau) indicates tau pathology and correlates with cognitive decline, while total tau (t-Tau) marks ongoing neuronal injury and axonal degeneration ^1,4,5^. The Aβ42/p-Tau ratio integrates both pathologies and demonstrates superior diagnostic accuracy by capturing the balance between amyloid burden and tau-mediated neurodegeneration ^1,4,5^. Despite their accuracy, CSF biomarkers require invasive lumbar puncture creating significant screening barriers. As a valuable alternative, blood-based biomarkers have emerged in recent years as a non-invasive alternative for AD diagnosis, with phosphorylated tau at threonine 217 (plasma p-Tau217) providing high sensitivity and specificity for AD diagnosis ^6^. However, these biochemical markers while detecting molecular pathology, these kind of biomarkers provide no information about functional status of neural networks affected by these processes, particularly attention networks vulnerable to early pathology ^7^. This gap is clinically significant because individuals can exhibit molecular pathology without corresponding functional network impairment, or conversely, show network dysfunction with varying degrees of molecular burden. Understanding this relationship is essential for predicting symptom onset and monitoring disease progression.

The relationship between molecular pathology and functional network disruption involves distinct but interacting mechanisms. Post-mortem studies identify the locus coeruleus (LC) as one of the earliest sites of tau accumulation, appearing before cortical involvement ^8^. This brainstem nucleus serves as the brain’s primary noradrenergic center with extensive projections to cortical attention, arousal, and executive networks ^9,10^. Simultaneously, cortical amyloid accumulation follows consistent spatiotemporal ordering in regions overlapping the default mode network (DMN), including posterior cingulate cortex, precuneus, and medial orbitofrontal cortex ^11,12^. Critically, elevated amyloid burden in cognitively healthy individuals associates with failure of DMN suppression during goal-directed tasks ^11,12^. This persistent DMN activity creates interference with task-positive networks, including attention systems, resulting in impaired performance and accelerated decline ^12,13^. Thus, the LC’s vulnerability to tau pathology, combined with amyloid-driven DMN dysfunction, creates dual mechanisms of attentional network disruption that necessitate assessment tools capable of capturing both LC-mediated attention engagement and DMN suppression during cognitively demanding tasks.

The oddball paradigm provides an ideal framework for examining these LC-attention-oculomotor interactions through a well-characterized temporal cascade ^14^. Infrequent target stimuli embedded among frequent distractors robustly activate the LC, initiating coordinated activity across attention networks ^9,15^: the ventral attention network (VAN) responds preferentially to salient oddballs, the dorsal attention network (DAN) coordinates top-down attentional orienting ^16^, and the frontoparietal network (FPN) supports sustained executive control ^17,18^. Critically, successful performance requires DMN suppression, as failure to deactivate DMN regions characterizes individuals with elevated amyloid ^19,20^. This temporal progression from LC activation through attention engagement with concurrent DMN suppression creates sensitivity for detecting functional alterations from both LC tau pathology and amyloid-driven DMN dysfunction.

The anatomical connections between LC and oculomotor control systems may enable non-invasive capture of this temporal cascade. The LC projects directly to the Edinger-Westphal nucleus, making pupil dynamics a proxy for LC phasic firing, with larger dilations for task-relevant stimuli reflecting noradrenergic modulation of arousal and attention ^21,22^. Additionally, the LC modulates the superior colliculus, coordinating binocular vergence movements that correlate with visual attention allocation and reflect late-component event-related potentials in parietal attention networks ^23,24^. The vergence-pupil interaction follows a well-documented temporal sequence, with cognitive vergence movements preceding and conditioning subsequent pupillary changes ^24,25^. This temporal pattern, combined with LC modulation and cortical oculomotor responses, suggests that cognitive vergence and pupil dynamics could capture activity across attention networks ^21–23^. Previous evidence demonstrates altered cognitive vergence and pupil responses during oddball tasks in MCI and AD patients ^24^. Additionally, saccadic eye movements and pupil dilation have been identified as potential indicators of MCI and AD ^26,27^. This suggest that these measures may be sensitive to functional consequences of pathology affecting LC-attention circuits.

However, the direct relationship between CSF biomarker levels and attention-dependent oculomotor responses remains poorly characterized. Given that molecular pathology affects the neural substrates supporting attentional differentiation, tau accumulating in the LC and amyloid disrupting DMN suppression ^5,8^, altered CSF biomarker levels may modulate attentional performance as operationalized in the oddball task through diminished capacity to differentiate targets from distractors via cognitive vergence and pupil responses. Given that cognitive vergence and pupil dynamics reflect activity within the attentional networks, pathological accumulation of p-Tau or amyloid could manifest as altered oculomotor discrimination patterns. Given the shared embryological origin of ocular and neural tissues ^28^, establishing these relationships would reveal how molecular pathology translates into measurable functional alterations, supporting oculomotor assessment as an accessible functional biomarker that complements biochemical measures and enables detection of attention network dysfunction before overt cognitive symptoms, while clarifying the relative contributions of tau and amyloid pathology.

Despite evidence of oculomotor alterations in AD, critical gaps remain in characterizing how molecular pathology relates to functional network capacity. Most studies have examined resting-state or trait-based oculomotor metrics rather than task-evoked responses during cognitively demanding paradigms that specifically engage LC-attention circuits (26,27). Task-based approaches capturing stimulus differentiation capacity are mechanistically more informative because they index dynamic attention network engagement and DMN suppression under cognitive control demands—the networks vulnerable to tau and amyloid pathology. Moreover, existing studies have relied primarily on categorical comparisons (AD versus controls) rather than examining continuous relationships between quantitative CSF biomarker levels and functional oculomotor performance. To our knowledge, no studies have directly examined how CSF biomarker concentrations relate to oculomotor stimulus differentiation during attention tasks, limiting understanding of how specific molecular pathologies translate into measurable functional alterations and which pathological process most strongly associates with attentional control disruption.

Therefore, this study aimed to determine whether CSF biomarkers (Aβ42, p-Tau, t-Tau, and Aβ42/p-Tau ratio) are associated with altered stimulus differentiation in cognitive vergence and pupil responses during an oddball task, and to evaluate oculomotor metrics as predictors of CSF core AD biomarkers in patients at MCI stage.

## 2. Methods

This study employed a cross-sectional design. The study was approved by the Ethics Committees of the University of Barcelona (HCB/2021/0668) and Hospital del Mar. All methods were performed in accordance with the relevant institutional and national guidelines and regulations. The research was conducted in accordance with the STROBE guidelines for cross-sectional studies ^29^, and the checklist is available in the supplementary material.

### 2.1 Participants

A total of 38 participants (12 men [31.6%] and 26 women [68.4%]) were included in the study. All participants had been clinically assessed at two hospitals in Barcelona, Spain (Hospital del Mar and Hospital Clínic de Barcelona), where CSF collection via lumbar puncture was performed as part of routine clinical workup. Based on these clinical assessments, all participants showed abnormal CSF core AD biomarkers in accordance with Alzheimer’s Association criteria ^30^, placing them within the Alzheimer’s disease biological continuum at the MCI clinical stage. Consistent with the NIA-AA biological framework, the MCI diagnosis reflects the current clinical syndrome independently of the underlying AD molecular pathology ^30^.

Participants and their families received detailed instructions about the study procedures. Before entering the study, patients or relatives signed a written informed consent for their participation in accordance with the Helsinki Declaration.

### 2.2 Sample Size

Analyses included 38 participants drawn from a larger study cohort, with enrollment determined by availability of CSF biomarker data (a convenience subsample). Because sample size was determined by biomarker availability rather than formal a priori power analysis, we conducted post-hoc power analysis to characterize the study’s sensitivity to detect the observed effects.

Power analysis for linear mixed-effects models with hierarchical time-series data is complex and cannot be adequately addressed using standard analytical formulas, which assume independent observations. We therefore employed simulation-based methods to estimate power, which appropriately account for the nested data structure (time points within participants) and the specific random effects structure of our models (random intercepts and slopes for time).

For the model showing the strongest interaction effect (Vergence ∼ CSF p-Tau), we simulated 1,000 datasets based on the fitted model parameters and re-estimated the mixed-effects model for each simulated dataset. With n=38 participants and the observed interaction effect (β = -0.035, SE = 0.0047, p = 3.8×10□^1^ □), the study achieved power >99% for detecting this effect at α = 0.05. All 1,000 simulated datasets yielded significant interactions (p < 0.05), with 100% yielding p < 0.001. This high power reflects both the magnitude of the observed effect and the precision gained from averaging across approximately 6,800 temporal observations per participant.

To assess sensitivity to sample size, we conducted power curve analysis by randomly subsampling participants and re-estimating power. Results indicated that power remains >99% with n≥25 participants but drops substantially to 56% at n=20. Results indicated that power remains >99% with n≥25 participants, but drops substantially to 56% at n=20, confirming the validity of the simulation approach and indicating adequate sensitivity for detecting the observed interaction effects.

### 2.3 Inclusion/Exclusion Criteria

#### Inclusion criteria

Participants were enrolled if they: (1) were aged 65 years or older; (2) had undergone CSF biomarker assessment (Aβ42, p-Tau, t-Tau) via lumbar puncture during routine clinical workup; (3) possessed sufficient visual function (corrected or uncorrected) to discriminate on-screen stimuli; and (4) were able to provide written informed consent, either personally or through a legally authorized representative.

#### Exclusion criteria

Participants were excluded for: (1) severe cognitive impairment (Mini-Mental State Examination <10 ^31^); (2) neurological disorders with documented cognitive consequences beyond MCI/AD (e.g., Parkinson’s disease); (3) major psychiatric illness; (4) structural brain lesions on neuroimaging contributing to cognitive deficits (e.g., intracranial masses); (5) ophthalmologic pathology precluding task completion (blindness, strabismus, nystagmus, or significant retinal/oculomotor disease); and (6) insufficient Spanish proficiency (oral or written).

### 2.4 Device

Visual stimuli were presented and oculomotor data were recorded using the BGaze system (BraingazeSL, Mataró, Spain). Eye position and pupil diameter were captured with a remote eye tracker (Tobii 5L; 33 Hz; Tobii Technology AB, Sweden). According to the manufacturer’s specifications, the system achieves a binocular gaze accuracy of 0.4° and a precision of 0.32° of visual angle under optimal viewing conditions.

### 2.5 CSF Core AD Biomarkers

Aβ42, tTau, and pTau were determined using the single molecule array technology implemented on the Lumipulse platform, in accordance with standardized analytical procedures ^5^. Furthermore, the Aβ42/pTau ratio was computed to provide an additional indicator of neurodegeneration. All biochemical assessments were performed by certified professionals at the participants’ respective hospital laboratories.

### 2.6 Procedure

Testing took place in a dimly lit room with curtains drawn and overhead lights off. Participants were seated at approximately 50 cm from the display screen, with the eye tracker positioned below it. Use of corrective lenses was permitted. A chinrest was employed throughout the session to minimize head displacement.

### 2.7 Paradigm

A visual oddball paradigm was administered, following the protocol described by Jiménez et al. ^24^ with minor adaptations. An oddball paradigm was applied. An oddball task activates many brain regions and is used to assess neural activity in the LC ^15^. The task comprised 120 trials. Each trial began with a 2000 ms grey mask screen, followed by a centrally presented letter string (11 characters, upper or lower case, randomly selected) displayed for 2000 ms (Figure 1). Letter strings were semantically meaningless and did not form acronyms or words. The only dimension differentiating conditions was font color: blue strings constituted the frequent distractor condition (80% of trials) and red strings the infrequent target condition (20% of trials). Participants were instructed to fixate on the screen and press a response button exclusively upon detecting red stimuli. Verbal reminders were provided by the experimenter when necessary. Trial order was randomized. Binocular eye movements were recorded continuously, and the tracker was calibrated using a five-point binocular procedure prior to task onset. Total task duration was approximately six minutes.

**Figure 1.**
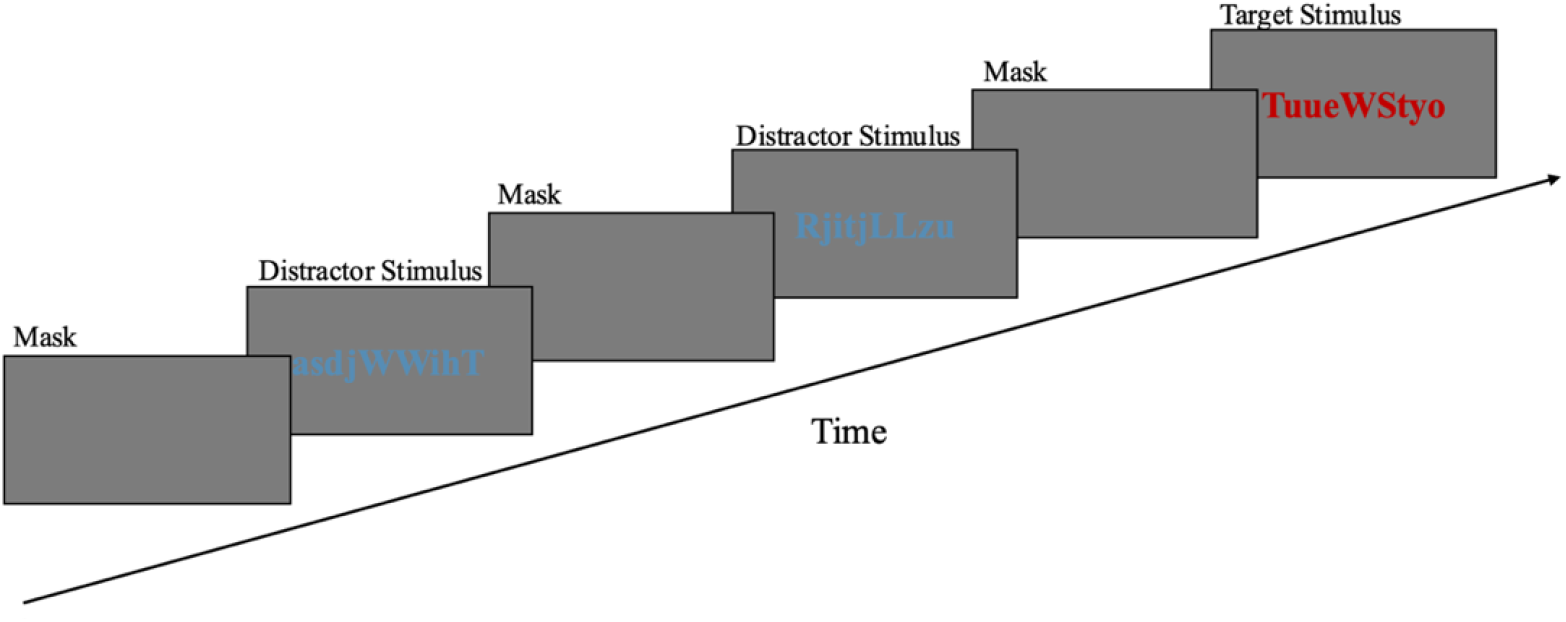
Oddball Task sequence. Strings of symbols are presented for two seconds, followed by a grey mask. In 80% of the trials the symbols were printed in blue, and in the remaining 20% they were printed in red.

### 2.8 Statistical Analysis

First, eye-tracking data were preprocessed in several sequential steps. First, outlier rejection was applied: fixations with normalized screen coordinates below zero (off-screen) and vergence values outside the physiologically plausible range (0°–20°) were discarded. Trials with more than 30% invalid samples were excluded. For retained trials, gaps in the signal were reconstructed using linear interpolation, following established recommendations ^32^. After excluding trials with more than 30% missing samples, the remaining trials showed limited need for reconstruction: 71.8% required ≤20% interpolation and 28.2% required <5%, consistent with expected blink-related signal interruptions in a clinical population.

Second, screen pixel coordinates were converted to physical units based on display dimensions (340 mm × 190 mm). Vergence angles were derived using a vector-based geometric approach ^23^: at each sample, unit directional vectors were computed from each eye’s position to the fixation target, and the vergence angle θ was obtained via the dot product of these vectors, expressed in degrees. Baseline-corrected vergence was computed as υ(t) = γ(t) − γ□. For the pupillometric signal, binocular diameter values were averaged across eyes at each time point, and baseline-corrected pupil responses were obtained as π(t) = p(t) − p□. The baseline reference (γ□, p□) was defined as the mean signal over the first seven samples (∼212 ms pre-stimulus), in line with recommended practice ^32^. Both signals were smoothed with a Gaussian moving-average filter prior to baseline correction. Condition-level response curves were then obtained by averaging υ(t) and π(t) across trials within each condition (target, distractor).

Third, to avoid assuming linearity, we first explored biomarker-oculomotor associations using Generalized Additive Mixed Models, which can detect complex non-linear relationships. All models demonstrated approximately linear relationships (effective degrees of freedom = 1.01-1.07), confirming the appropriateness of linear modeling for our data. Therefore, we examined whether CSF biomarkers predicted differential oculomotor responses to target versus distractor stimuli using linear mixed-effects models. Vergence and pupil time series were modeled using linear mixed-effects models including fixed effects of Condition, Biomarker, their interaction, and covariates (Age, Sex and MMSE), with random intercepts and participant-specific random slopes for Time to account for individual temporal trajectories. The critical Condition × Biomarker interaction tested whether the magnitude of stimulus differentiation varied as a function of biomarker level. All continuous predictors and outcomes were standardized (Z-scored) prior to analysis.

Because the data consist of high-resolution temporal sequences nested within participants, traditional multiple comparison corrections such as Bonferroni or FDR are not appropriate, as they assume independent observations. Instead, we adopted the principle of “keep it maximal”, in order to prevent type I error, including the necessary random effects (here, participant and the slope of time) to appropriately model the hierarchical structure and temporal dependencies ^33^. Time was included both as a random slope (to capture participant-specific temporal dynamics) and as a covariate, which allows estimation of the average temporal trend while separating it from individual deviations, improving model interpretability and controlling for within-trial temporal effects.

Our hierarchical approach included: a) Models using the full vergence and pupil time series as dependent variables. b) Decomposition into temporal and magnitude features. For biomarkers showing significant Condition × Biomarker interactions at level (a), we further analyzed trial-level features (100 trials per participant), including temporal/shape features (initial slope, global slope, late slope, time to peak) and magnitude features (AUC, peak amplitude). At this level, time was not included as a random slope, because collapsing the data to trial-level averages removes within-trial temporal resolution. Each feature was modeled using the same mixed-effects specification to identify the mechanistic source of response modulation. The preprocessing pipeline (linear interpolation of short gaps and Gaussian smoothing) ensured temporal continuity and reduced high-frequency noise, enabling stable estimation of slope- and peak-based features without altering the underlying response dynamics. c) Reverse prediction: ocular features as predictors of CSF biomarkers. Features with significant Condition × Biomarker interactions were aggregated at the participant level and tested as predictors of biomarker concentrations using linear regression to explore the diagnostic potential of oculomotor differentiation metrics with Bonferroni correction. Given the exploratory nature of this analysis and relative to sample size (N=38), cross-validation was not employed. The univariate regression approach with single predictors minimizes overfitting risk, and our focus was on identifying potential biomarker-oculomotor relationships rather than developing a validated diagnostic tool.

To improve interpretability, all values (vergence, pupil, trial-level features, and biomarkers) were standardized (Z-scored). Residual distributions were examined via histograms and Q-Q plots. Residuals from pupil models were approximately normal, with smooth tails. For vergence models, histograms suggested normality, though Q-Q plots showed slight tail deviations. To assess robustness, we compared Gaussian linear mixed models against generalized linear mixed models with a Student’s t error distribution, which are less sensitive to heavy tails. Gaussian models yielded lower AIC and BIC (Gaussian: AIC = 897.495, BIC = 897.579 vs. Student’s t: AIC = 898.196, BIC = 898.279), supporting their use. This approach also avoids data transformations that could alter the fundamental logic of eye movement data (positive = convergence, negative = divergence). Non-normality is not a major concern given the large dataset, as linear mixed moles are robust to moderate deviations from normality in larger time series samples. All models were adjusted for sex, age, and MMSE scores, with significance set at p < 0.05. Analyses were conducted in Python 3.x (pandas, numpy, scipy, matplotlib, statsmodels).

## 3. Results

Table 1 summarizes participant characteristics. The sample showed considerable variability in CSF biomarker levels, particularly in p-Tau and t-Tau, reflected in the wide distribution of Aβ42/p-Tau ratios. This heterogeneity enabled examination of continuous biomarker-oculomotor relationships across a broad pathological spectrum. Average oculomotor responses (Figure 2) demonstrated preserved stimulus differentiation at the group level: targets elicited enhanced vergence convergence and sustained pupillary dilation compared to distractors, with considerable interindividual variability (shaded CI 95% bands)

**Table 1.** Participants Characterization.

| Variable | Mean ± SD |
| --- | --- |
| Age (y) | 72.9 ± 5.83 |
| MMSE | 22.4 ± 4.04 |
| Aβ42 | 623 ± 281 |
| p-Tau | 160 ± 196 |
| t-Tau | 762 ± 370 |
| Aβ42/p-Tau ratio | 7.80 ± 7.77 |

y: years. MMSE: Mini- Mental State Examination.

**Figure 2.**
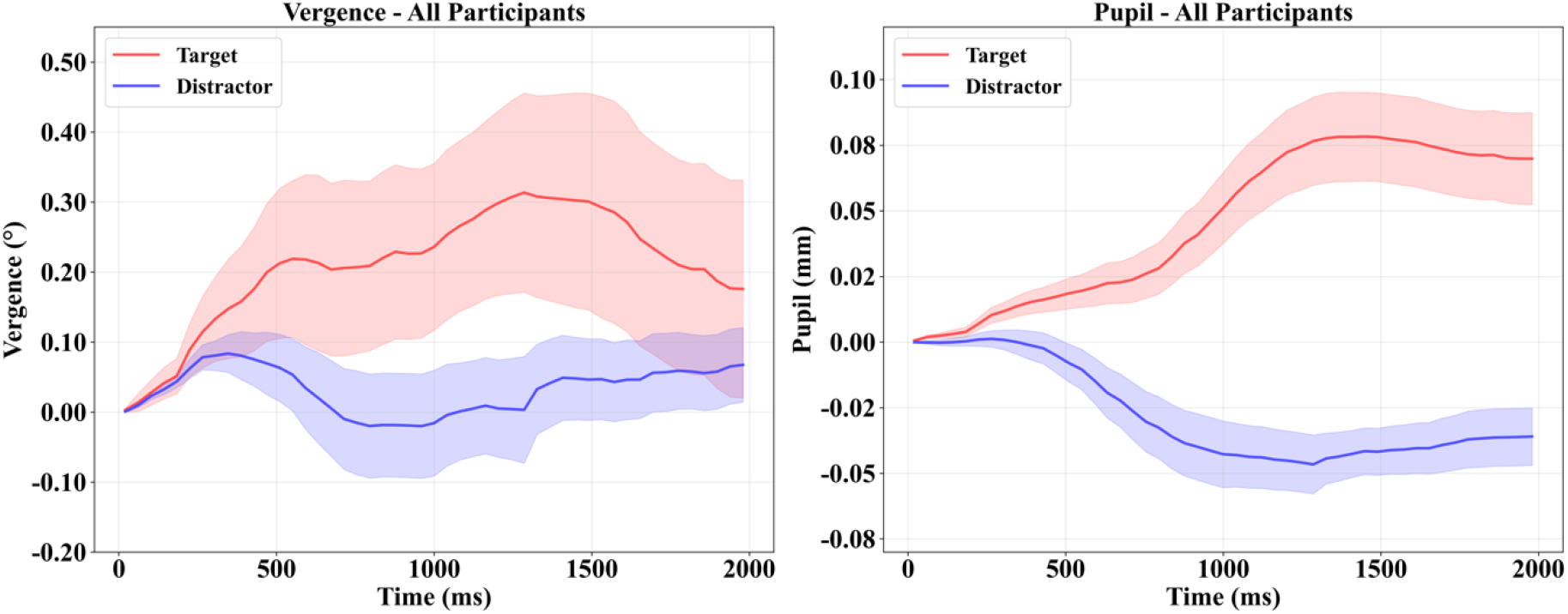
Cognitive vergence and pupil main curves Vergences in degrees and pupil in millimeters. Shaded regions: CI95%

### 3.1 Full vergence and pupil responses

The target vs. distractor difference was significant across all vergence and pupil models (Vergence: β ≈ 0.066, SE ≈ 0.00479, p < 0.001; Pupil: β ≈ 0.451, SE ≈ 0.00465, p < 0.001). Main effects of biomarkers were not significant in any model. However, condition × biomarker interactions revealed that biomarker levels were systematically associated with stimulus differentiation (Figure 3). Specifically, higher Aβ42 and Aβ42/p-Tau levels showed positive associations with target-distractor differentiation in vergence (Aβ42: β = 0.013, SE = 0.00482, p = 0.005; Aβ42/p-Tau: β = 0.029, SE = 0.00479, p < 0.001), while Aβ42 showed a negative association with pupil differentiation (Aβ42: β = -0.019, SE = 0.00468, p < 0.001) and Aβ42/p-Tau showed a positive association (β = 0.011, SE = 0.00465, p = 0.015). Higher CSF p-Tau levels were negatively associated with differentiation in both vergence (β = -0.035, SE = 0.00470, p < 0.001) and pupil responses (β = -0.060, SE = 0.00456, p < 0.001). CSF t-Tau showed no significant associations with oculomotor differentiation.

**Figure 3.**
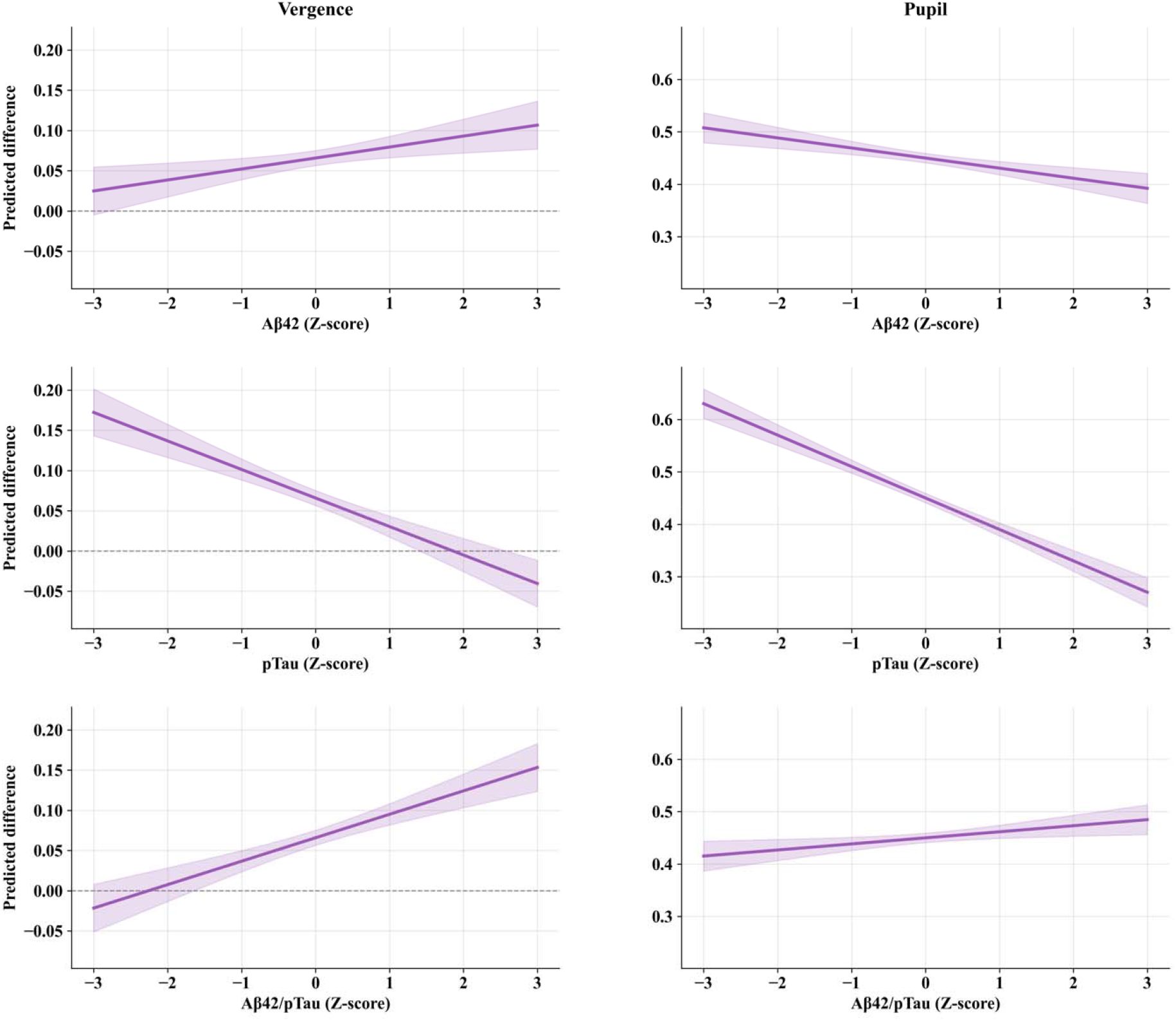
Condition × biomarker interaction effects on oculomotor differentiation. For vergence (left) and pupil (right) responses. Lines show how the magnitude of the condition effect (target vs. distractor) scales with biomarker concentration. Positive slopes indicate that higher biomarker levels amplify differentiation (Aβ42, Aβ42/pTau in vergence); negative slopes indicate reduced differentiation (pTau in both measures; Aβ42 in pupil). Shaded regions: CI95%. Ocular response and CSF biomarkers are Z-score standardized.

### 3.2 Features response

From the complete vergence and pupil responses, we extracted temporal/shape features (initial slope, global slope, late slope, time to peak) and magnitude features (AUC, peak amplitude) for Aβ42, p-Tau, and the Aβ42/p-Tau ratio, which showed significant interactions in the full response models. Several biomarker main effects and interactions were identified in relation to oculomotor response features. For vergence features, CSF p-Tau showed a main association with Slope Initial, while condition × biomarker interactions were observed between CSF Aβ42 and Time to Peak, p-Tau and Time to Peak, and p-Tau and Peak Power. For pupil features, CSF p-Tau showed a main association with Slope Global. Regarding interactions, higher CSF p-Tau levels were associated with reduced target-distractor differences in Peak Power and Time to Peak (negative β values for interactions), consistent with diminished pupillary differentiation at higher tau burden. In contrast, higher Aβ42/p-Tau ratios were associated with enhanced differences in Peak Power and Time to Peak (positive β values), consistent with preserved pupillary discrimination capacity in more favorable pathological profiles. The interaction in Slope Late with the Aβ42/p-Tau ratio showed that higher ratios were associated with reduced slope differences between conditions. These significant findings are detailed in Table 2; the full results are available in supplementary material.

**Table 2.** Features results.

| Feature | Biomarker | Effect | $\beta$ | SE | p |
| --- | --- | --- | --- | --- | --- |
| Vergence |  |  |  |  |  |
| Slope Initial | pTau | Condition | 0.024 | 0.0396 | 0.539 |
|  |  | Biomarker | -0.057 | 0.0257 | **0.025** |
|  |  | Interaction | -0.036 | 0.0388 | 0.349 |
| Time to Peak | Aβ42 | Condition | -0.007 | 0.0388 | 0.993 |
|  |  | Biomarker | -0.053 | 0.0470 | 0.257 |
|  |  | Interaction | 0.108 | 0.0390 | **0.005** |
| Time to Peak | pTau | Condition | -0.001 | 0.0388 | 0.998 |
|  |  | Biomarker | 0.039 | 0.0461 | 0.395 |
|  |  | Interaction | -0.087 | 0.0381 | **0.022** |
| Peak Power | pTau | Condition | 0.011 | 0.0353 | 0.746 |
|  |  | Biomarker | -0.017 | 0.0803 | 0.832 |
|  |  | Interaction | -0.085 | 0.0347 | **0.013** |
| Pupil |  |  |  |  |  |
| Slope Global | pTau | Condition | 0.724 | 0.0358 | **<0.001** |
|  |  | Biomarker | -0.130 | 0.0528 | **0.013** |
|  |  | Interaction | -0.072 | 0.0352 | **0.040** |
| Slope Late | Aβ42/pTau | Condition | -0.044 | 0.0396 | 0.264 |
|  |  | Biomarker | -0.015 | 0.0256 | 0.555 |
|  |  | Interaction | -0.091 | 0.0396 | **0.020** |
| AUC | pTau | Condition | 0.560 | 0.0370 | <0.001 |
|  |  | Biomarker | -0.068 | 0.0519 | 0.188 |
|  |  | Interaction | -0.073 | 0.0364 | **0.043** |
| Time to Peak | pTau | Condition | 0.587 | 0.0370 | **<0.001** |
|  |  | Biomarker | -0.042 | 0.0499 | 0.391 |
|  |  | Interaction | -0.128 | 0.0364 | **<0.001** |
| Time to Peak | Aβ42/pTau | Condition | 0.586 | 0.0371 | **<0.001** |
|  |  | Biomarker | 0.053 | 0.0486 | 0.269 |
|  |  | Interaction | 0.072 | 0.0370 | **0.049** |
| Peak Power | pTau | Condition | 0.592 | 0.0362 | **<0.001** |
|  |  | Biomarker | 0.021 | 0.0580 | 0.712 |
|  |  | Interaction | -0.239 | 0.0356 | **<0.001** |
| Peak Power | Aβ42/pTau | Condition | 0.591 | 0.0364 | **<0.001** |
|  |  | Biomarker | -0.013 | 0.0567 | 0.817 |
|  |  | Interaction | 0.115 | 0.0364 | **0.002** |

All models included sex, MMSE and age as covariates. $\beta$ represents standardized regression coefficients. SE = standard error. Condition effect represents the difference between Target and Distractor trials. Biomarker effect represents the main effect of the respective CSF biomarker (Aβ42, pTau, or Aβ42/pTau ratio). Interaction represents the Condition × Biomarker interaction term.

### 3.3 Ocular response as biomarker predictor

Features were averaged at the participant level and included as predictors in linear regression models with interaction terms, where biomarkers served as the dependent variable. The models tested main effects, condition effects, and feature × condition interactions. While interactions were not statistically significant, two main effects reached significance, both involving pTau: vergence slope initial showed a significant negative association (β = -0.784, SE = 0.315, p = 0.015, R^2^ = 0.20), and pupil slope global also demonstrated a significant negative relationship (β = -0.610, SE = 0.277, p = 0.016, R^2^ = 0.21). The remaining regressions are reported in the supplementary material.

## 4. Discussion

This study aimed to determine whether CSF biomarkers (Aβ42, p-Tau, t-Tau, and Aβ42/p-Tau ratio) are associated with altered stimulus differentiation in cognitive vergence and pupil responses during an oddball task, and to evaluate oculomotor metrics as predictors of CSF core AD biomarkers in patients at MCI stage. Our findings suggest that CSF p-Tau is consistently associated with reduced oculomotor differentiation (distractor vs target), while Aβ42 shows divergent associations with cognitive vergence versus pupil systems. These associations manifested in temporal slope features, magnitude, and timing metrics. Additionally, oculomotor features from showed predictive associations with CSF p-Tau levels in reverse prediction analyses.

Our results show a consistent pattern in which higher CSF p-Tau levels were negatively associated with oculomotor differentiation capacity, reflected in reduced vergence and pupillary dilation metrics, with the most pronounced associations observed for pupillary power. This pattern is consistent with evidence that the LC is one of the earliest sites of tau accumulation ^8^, which is associated with altered noradrenergic modulation involved in inhibitory control, attentional orientation, and arousal regulation ^10,15^. LC pathology is associated with compromised filtering of irrelevant information and reduced response adjustment to behaviorally relevant stimuli, as the LC modulates cortical synchronization through alpha and theta waves and influences activation of attentional networks (DAN, VAN, and FPN) ^16–20,35^. These converging mechanisms are consistent with our observation that individuals with greater CSF p-Tau burden showed reduced differentiation in both vergence and pupillary responses. The concurrent impairment in both measures may reflect a shared pathway, as vergence movements directly modulate pupillary responses through established vergence-pupil coupling mechanisms ^21,25^. Thus, tau-associated impairment in vergence control may secondarily influence pupillary differentiation through this coupling.

In contrast, higher CSF Aβ42 levels, indicative of lower amyloid pathology, were associated with greater vergence differentiation but reduced pupillary differentiation. Importantly, in this context, reduced pupillary differentiation may reflect greater processing efficiency rather than impaired function. This pattern closely resembles the response profile observed in cognitively normal controls in prior literature ^24^: healthy individuals demonstrated robust vergence-based target detection with relatively modest pupillary responses, whereas MCI/AD patients showed the opposite—reduced vergence differentiation accompanied by exaggerated pupillary responses. The current findings demonstrate that within this clinical cohort, the degree of preservation of this ‘healthy’ response profile scales continuously with Aβ42 levels, providing a functional readout of relative network integrity.

Higher Aβ42 levels may be associated with preserved functional integrity of oculomotor and attentional networks involving the VAN, DAN, and FPN, which integrate structures such as the superior colliculus, frontal eye fields, and parietal cortex ^12,16^. This preserved network integrity was associated with more precise oculomotor modulation during target detection, as reflected by greater vergence differentiation. Concurrently, intact attentional networks may be associated with reduced need for compensatory arousal mechanisms, resulting in lower pupillary differentiation. Previous studies have demonstrated that individuals with MCI/AD exhibit hyperreactive pupillary responses, interpreted as reflecting compensatory increases in arousal ^24,27^. This hyperreactivity may arise from amyloid-related disruption of DMN suppression during goal-directed tasks, which interferes with attentional networks and is associated with greater dependence on compensatory arousal mechanisms ^12,36^. When the DMN fails to deactivate properly, attentional efficiency is compromised, manifesting in greater vergence divergence toward relevant stimuli that may condition exaggerated pupillary dilation through the vergence-pupil coupling mechanism ^12,25,36–38^.

Therefore, individuals with more preserved Aβ42 levels may require less compensatory arousal modulation, maintaining preserved vergence control while exhibiting reduced pupillary differentiation. Future studies should confirm this interpretation with concurrent neuroimaging of DMN suppression and autonomic measures in larger and more diverse samples.

The Aβ42/p-Tau ratio showed positive associations with differentiation in both vergence and pupillary responses. This pattern is consistent with the interpretation that this ratio reflects functional reserve, capturing the balance between amyloid-induced network disruption and tau-mediated neurodegeneration. This interpretation aligns with evidence demonstrating that the ratio provides superior diagnostic value compared to individual biomarkers ^4,39,40^. More favorable ratios were associated with relative preservation of attentional circuits and arousal systems, consistent with operation within a non-compensatory regime. The observed pattern of reduced pupillary differentiation in late phases aligns with processing efficiency principles and the adaptive gain theory of LC function ^41^. Optimal norepinephrine activity during early phases is associated with enhanced signal-to-noise ratios for target processing ^42^, following the Yerkes-Dodson law’s inverted-U relationship between arousal and performance. Following successful early target detection, prolonged arousal modulation may become unnecessary, and the locus coeruleus-norepinephrine system may adaptively downregulate, manifesting as reduced pupillary responses in late phases. This pattern aligns with studies linking reduced sustained pupillary dilation with more efficient and less cognitively costly processing ^43^.

Reverse prediction analyses showed that oculomotor features contain quantifiable information about molecular pathology. Features were averaged at the participant level and included as predictors in linear regression models with biomarkers as the dependent variable. While feature × condition interactions were not statistically significant, main effects of oculomotor slope features showed significant associations with CSF p-Tau levels: both vergence slope initial and pupil slope global were negatively associated with CSF p-Tau. No significant associations were observed with Aβ42. This selective association with CSF p-Tau has a mechanistic basis: the inhibition of responses to irrelevant stimuli critically depends on noradrenergic control from the LC ^15,38,41^, and this nucleus is a primary early site of tau accumulation, even before detectable cortical involvement ^8^. Oculomotor slope features, which reflect the temporal dynamics of response initiation and modulation, may be particularly sensitive to tau-associated LC alterations affecting noradrenergic fine-tuning ^15,21^. If validated in larger samples, such metrics could complement biochemical assays in clinical monitoring contexts where repeated CSF sampling is impractical.

Therefore, oculomotor responses during distractor processing, which require inhibitory control and fine-tuning of noradrenergic tone, may be particularly sensitive to tau-associated LC alterations ^15,35^. These findings reinforce the notion that oculomotor dynamics provide a sensitive readout of LC integrity, capturing functional consequences of tau pathology before widespread cortical involvement, and highlighting the sequential vulnerability of attention and oculomotor circuits in the progression of AD.

### 4.1 Clinical implications

These findings suggest that oculomotor differentiation metrics extracted during attention tasks may serve as accessible functional biomarkers complementing biochemical CSF and blood-based measures. While CSF biomarkers detect molecular pathology and are associated with cognitive decline trajectories ^5,6^, they provide limited information about the real-time functional status of attentional networks. In contrast, oculomotor measures may capture the dynamic capacity of LC-attention circuits to process salient information, potentially bridging the gap between molecular pathology and clinical symptoms. Eye-tracking technology is non-invasive, cost-effective, and scalable, making it suitable for repeated assessments in clinical monitoring or therapeutic trials targeting attention and arousal systems.

The specificity of CSF p-Tau associations with oculomotor differentiation suggests these metrics could serve as functional readouts of tau-associated LC alterations, complementing plasma biomarkers such as p-Tau217 ^6^. Taken together, these findings provide preliminary evidence that temporal oculomotor dynamics may capture functional signatures associated with tau burden, potentially with greater specificity than DMN network alterations associated with amyloid. These metrics represent promising candidates for functional biomarkers of LC-related dysfunction and, due to their specificity to CSF p-Tau, as potential tools for characterizing biological profiles. Future work should validate these metrics in larger and preclinical populations, establish longitudinal relationships with symptom onset, and integrate them with neuroimaging and electroencephalography measures of LC integrity and network connectivity to test mechanistic hypotheses linking molecular pathology to functional circuit alterations.

### 4.2 Strengths and Limitations

This study has several strengths, to the best of our knowledge, this is the first study to examine continuous relationships between quantitative CSF biomarker levels and attention-dependent oculomotor differentiation, utilizing temporal feature extraction to localize pathology effects to specific processing phases, and employing bidirectional analyses to establish proof-of-principle that oculomotor dynamics contain information about molecular pathology. However, limitations include the modest sample size (n=38) that constrained statistical power particularly for reverse prediction analyses, the cross-sectional design that precludes determination of temporal precedence between oculomotor decline and symptom onset, restriction to MCI/AD populations limiting generalizability to preclinical stages where functional biomarkers might provide greatest clinical utility, absence of cognitively normal controls, and lack of concurrent neuroimaging to directly measure LC structural integrity or functional connectivity. Given these limitations, our results should be considered an initial proof-of-concept. Future work should validate these metrics in larger and preclinical populations, establish longitudinal relationships with symptom onset, and integrate them with neuroimaging and electroencephalography measures of LC integrity and network connectivity to test mechanistic hypotheses linking molecular pathology to functional circuit disruption.

## 5. Conclusion

This study demonstrates that higher CSF p-Tau levels are consistently associated with reduced oculomotor differentiation during attention tasks, while Aβ42 and Aβ42/p-Tau ratios show divergent associations with vergence versus pupil systems. These associations manifested in temporal slope features, magnitude, and timing metrics, suggesting that oculomotor dynamics may capture functional consequences of molecular pathology affecting LC-attention circuits. Oculomotor features showed predictive associations with CSF p-Tau levels, highlighting their potential sensitivity to tau-associated LC alterations. These findings highlight the potential of eye-tracking metrics as non-invasive tools that could complement biochemical CSF measures in identifying early attention network disruption in patients with MCI.

## Acknowledgment

R.M-F was supported by a grant from the National Agency for Research and Development (ANID)/Scholarship Program/DOCTORADO BECAS CHILE/2024–(Grant Nº 72240103. NF was the recipient of the Juan Rodés contract JR22/00014 (Instituto de Salud Carlos III, Spain). AI is supported by grants from the Multi-partner consortium to expand dementia research in Latin America [ReDLat, supported by Fogarty International Center (FIC), National Institutes of Health, National Institutes of Aging (R01 AG057234, R01 AG075775, R01 AG21051, R01 AG083799, CARDS-NIH, R01 AG057234), Alzheimer’s Association (SG-20-725707), Rainwater Charitable Foundation – The Bluefield project to cure FTD, and Global Brain Health Institute)], ANID/FONDECYT Regular (1250091 and 1210176 and 1220995); ANID/PIA/ANILLOS ACT210096; JPI JPND-Care, DISCeRN 2025 - Health and Social Care Research with a Focus on the Moderate and Late Stages of Neurodegenerative Diseases; FONDEF ID20I10152, and ANID/FONDAP 15150012; Wellcome Trust award for BRAIN-CLIMA: Investigating the Combined Impact of Heat and Air Pollution on Blood-Brain Barrier Integrity and Brain Aging in Latin America, (335293/Z/25/Z), and the CliCBrain (Horizon ID: 101236426; DOI 10.3030/101236426, Marie Skłodowska-Curie Actions - MSCA). The contents of this publication are solely the responsibility of the authors and do not represent the official views of these institutions. The funders had no role in study design, data collection and analysis, decision to publish or preparation of the manuscript.

## Author contributions

The corresponding author (H.S.) attests that all listed authors meet authorship criteria. HS and R.M-F they carried out the conceptualization of the study. HS secured and administered the funding that supported this study. R.M-F conducted statistical analysis. R.M-F and HS wrote the original draft. I.M-S, N.F, O. G-R, M S-C, C.C-M and A.I, performed a critical revision and editing of the manuscript. All authors have read and agreed to the published version of the manuscript.

## Data availability statement

Data is available via request to the corresponding author. The method of cognitive vergence for measuring neurodegenerative proteins has been protected by a patent application.

## Declaration of generative AI and AI-assisted technologies in the writing process

During the preparation of this work the author(s) used ChatGPT-4o in order to improve language and readability. After using this tool/service, the author(s) reviewed and edited the content as needed and take(s) full responsibility for the content of the publication.

## Patient consent statement

All participants provided informed written consent to participate in the study.

## References

1. Jack Jr., C. R. et al. NIA-AA Research Framework: Toward a biological definition of Alzheimer’s disease. Alzheimers Dement. 14, 535–562 (2018).

2. Prince, M. J. et al. World Alzheimer Report 2015 -The Global Impact of Dementia: An Analysis of Prevalence, Incidence, Cost and Trends. (Alzheimer’s Disease International, London, 2015).

3. GBD 2021 Nervous System Disorders Collaborators. Global, regional, and national burden of disorders affecting the nervous system, 1990-2021: a systematic analysis for the Global Burden of Disease Study 2021. Lancet Neurol. 23, 344–381 (2024).

4. Alcolea, D. et al. Amyloid precursor protein metabolism and inflammation markers in preclinical Alzheimer disease. Neurology 85, 626–633 (2015).

5. MilàUAlomà, M. et al. Amyloid beta, tau, synaptic, neurodegeneration, and glial biomarkers in the preclinical stage of the Alzheimer’s continuum. Alzheimers Dement. 16, 1358–1371 (2020).

6. Teunissen, C. E. et al. Blood-based biomarkers for Alzheimer’s disease: towards clinical implementation. Lancet Neurol. 21, 66–77 (2022).

7. Jiang, Y. et al. Alzheimer’s Biomarkers are Correlated with Brain Connectivity in Older Adults Differentially during Resting and Task States. Front. Aging Neurosci. 8, (2016).

8. Braak, H., Thal, D. R., Ghebremedhin, E. & Del Tredici, K. Stages of the pathologic process in Alzheimer disease: age categories from 1 to 100 years. J. Neuropathol. Exp. Neurol. 70, 960–969 (2011).

9. Dahl, M. J., Mather, M. & Werkle-Bergner, M. Noradrenergic modulation of rhythmic neural activity shapes selective attention. Trends Cogn. Sci. 26, 38–52 (2022).

10. Chandler, D. J., Gao, W.-J. & Waterhouse, B. D. Heterogeneous organization of the locus coeruleus projections to prefrontal and motor cortices. Proc. Natl. Acad. Sci. U. S. A. 111, 6816–6821 (2014).

11. Sperling, R. A. et al. The impact of Aβ and tau on prospective cognitive decline in older individuals. Ann. Neurol. 85, 181–193 (2019).

12. Anticevic, A. et al. The role of default network deactivation in cognition and disease. Trends Cogn. Sci. 16, 584–592 (2012).

13. Turner, G. R. & Spreng, R. N. Prefrontal Engagement and Reduced Default Network Suppression Co-occur and Are Dynamically Coupled in Older Adults: The Default–Executive Coupling Hypothesis of Aging. J. Cogn. Neurosci. 27, 2462–2476 (2015).

14. Murphy, P. R., O’Connell, R. G., O’Sullivan, M., Robertson, I. H. & Balsters, J. H. Pupil diameter covaries with BOLD activity in human locus coeruleus. Hum. Brain Mapp. 35, 4140–4154 (2014).

15. Krebs, R. M., Park, H. R. P., Bombeke, K. & Boehler, C. N. Modulation of locus coeruleus activity by novel oddball stimuli. Brain Imaging Behav. 12, 577–584 (2018).

16. Kim, H. Involvement of the dorsal and ventral attention networks in oddball stimulus processing: A meta-analysis. Hum. Brain Mapp. 35, 2265–2284 (2014).

17. Xiang, L. et al. Locus coeruleus noradrenergic neurons phase-lock to prefrontal and hippocampal infra-slow rhythms that synchronize to behavioral events. Front. Cell. Neurosci. 17, (2023).

18. Pahl, J. et al. Locus coeruleus integrity and left frontoparietal connectivity provide resilience against attentional decline in preclinical alzheimer’s disease. Alzheimers Res. Ther. 16, 119 (2024).

19. Raccah, O., Daitch, A. L., Kucyi, A. & Parvizi, J. Direct Cortical Recordings Suggest Temporal Order of Task-Evoked Responses in Human Dorsal Attention and Default Networks. J. Neurosci. Off. J. Soc. Neurosci. 38, 10305–10313 (2018).

20. Kucyi, A. et al. Electrophysiological dynamics of antagonistic brain networks reflect attentional fluctuations. Nat. Commun. 11, 325 (2020).

21. Joshi, S., Li, Y., Kalwani, R. M. & Gold, J. I. Relationships between Pupil Diameter and Neuronal Activity in the Locus Coeruleus, Colliculi, and Cingulate Cortex. Neuron 89, 221–234 (2016).

22. Joshi, S. & Gold, J. I. Pupil size as a window on neural substrates of cognition. Trends Cogn. Sci. 24, 466–480 (2020).

23. Sole Puig, M. et al. Attentional Selection Accompanied by Eye Vergence as Revealed by Event-Related Brain Potentials. PLoS ONE 11, e0167646 (2016).

24. Jiménez, E. C. et al. Altered Vergence Eye Movements and Pupil Response of Patients with Alzheimer’s Disease and Mild Cognitive Impairment During an Oddball Task. J. Alzheimers Dis. 82, 421–433 (2021).

25. Feil, M., Moser, B. & Abegg, M. The interaction of pupil response with the vergence system. Graefes Arch. Clin. Exp. Ophthalmol. 255, 2247–2253 (2017).

26. Opwonya, J. et al. Saccadic Eye Movement in Mild Cognitive Impairment and Alzheimer’s Disease: A Systematic Review and Meta-Analysis. Neuropsychol. Rev. 32, 193–227 (2022).

27. Granholm, E. L. et al. Pupillary Responses as a Biomarker of Early Risk for Alzheimer’s Disease. J. Alzheimers Dis. JAD 56, 1419–1428 (2017).

28. De Groef, L. & Cordeiro, M. F. Is the Eye an Extension of the Brain in Central Nervous System Disease? J. Ocul. Pharmacol. Ther. 34, 129–133 (2018).

29. von Elm, E. et al. The Strengthening the Reporting of Observational Studies in Epidemiology (STROBE) statement: guidelines for reporting observational studies. J. Clin. Epidemiol. 61, 344–349 (2008).

30. Jack Jr., C. R. et al. Revised criteria for diagnosis and staging of Alzheimer’s disease: Alzheimer’s Association Workgroup. Alzheimers Dement. 20, 5143–5169 (2024).

31. Tombaugh, T. N. & McIntyre, N. J. The mini-mental state examination: a comprehensive review. J. Am. Geriatr. Soc. 40, 922–935 (1992).

32. Mathôt, S. & Vilotijević, A. Methods in cognitive pupillometry: Design, preprocessing, and statistical analysis. Behav. Res. Methods 55, 3055–3077 (2023).

33. Barr, D. J., Levy, R., Scheepers, C. & Tily, H. J. Random effects structure for confirmatory hypothesis testing: Keep it maximal. J. Mem. Lang. 68, 255–278 (2013).

34. Schielzeth, H. et al. Robustness of linear mixed-effects models to violations of distributional assumptions. Methods Ecol. Evol. 11, 1141–1152 (2020).

35. Joshi, S. & Gold, J. I. Context-dependent relationships between locus coeruleus firing patterns and coordinated neural activity in the anterior cingulate cortex. eLife 11, e63490 (2022).

36. Wang, J. et al. Dysfunctional interactions between the default mode network and the dorsal attention network in subtypes of amnestic mild cognitive impairment. Aging 11, 9147–9166 (2019).

37. Fotiou, D. F. et al. Cholinergic deficiency in Alzheimer’s and Parkinson’s disease: evaluation with pupillometry. Int. J. Psychophysiol. Off. J. Int. Organ. Psychophysiol. 73, 143–149 (2009).

38. Murphy, P. R., Robertson, I. H., Balsters, J. H. & O’connell, R. G. Pupillometry and P3 index the locus coeruleus-noradrenergic arousal function in humans. Psychophysiology 48, 1532–1543 (2011).

39. Hansson, O. et al. CSF biomarkers of Alzheimer’s disease concord with amyloid-β PET and predict clinical progression: A study of fully automated immunoassays in BioFINDER and ADNI cohorts. Alzheimers Dement. J. Alzheimers Assoc. 14, 1470–1481 (2018).

40. Puig-Pijoan, A. et al. The CORCOBIA study: Cut-off points of Alzheimer’s disease CSF biomarkers in a clinical cohort. Neurologia 39, 756–765 (2024).

41. Aston-Jones, G. & Cohen, J. D. An integrative theory of locus coeruleus-norepinephrine function: adaptive gain and optimal performance. Annu. Rev. Neurosci. 28, 403#x2013;450 (2005).

42. Esterman, M. & Rothlein, D. Models of sustained attention. Curr. Opin. Psychol. 29, 174–180 (2019).

43. van der Wel, P. & van Steenbergen, H. Pupil dilation as an index of effort in cognitive control tasks: A review. Psychon. Bull. Rev. 25, 2005–2015 (2018).

