## Supplementary material for "Cognitive Vergence and Pupil Response During Oddball Task are Associated With Alzheimer’s Disease Cerebrospinal Fluid Neurodegenerative Biomarkers"

STROBE Statement—Checklist of items that should be included in reports of ***cross-sectional studies***

**Item**

|  | **No** | **Recommendation** | **N° page** |
| --- | --- | --- | --- |
| **Title and abstract** | 1 | (*a*) Indicate the study’s design with a commonly used term in the title or the abstract | 1-2 |
|  |  | (*b*) Provide in the abstract an informative and balanced summary of what was done  and what was found | 1-2 |
| **Introduction** |  |  |  |
| Background/rationale | 2 | Explain the scientific background and rationale for the investigation being reported | 2-4 |
| Objectives | 3 | State specific objectives, including any prespecified hypotheses | 4 |
| **Methods** |  |  |  |
| Study design | 4 | Present key elements of study design early in the paper | 4-7 |
| Setting | 5 | Describe the setting, locations, and relevant dates, including periods of recruitment,  exposure, follow-up, and data collection | 4-6 |
| Participants | 6 | (*a*) Give the eligibility criteria, and the sources and methods of selection of  participants | 4-5 |
| Variables | 7 | Clearly define all outcomes, exposures, predictors, potential confounders, and effect  modifiers. Give diagnostic criteria, if applicable | 5-7 |
| Data sources/ measurement | 8* | For each variable of interest, give sources of data and details of methods of assessment (measurement). Describe comparability of assessment methods if there is  more than one group | 5-7 |
| Bias | 9 | Describe any efforts to address potential sources of bias | 5-7 |
| Study size | 10 | Explain how the study size was arrived at | 5 |
| Quantitative variables | 11 | Explain how quantitative variables were handled in the analyses. If applicable,  describe which groupings were chosen and why | 5-7 |
| Statistical methods | 12 | (*a*) Describe all statistical methods, including those used to control for confounding | 7-8 |
|  |  | (*b*) Describe any methods used to examine subgroups and interactions | 7-8 |
|  |  | (*c*) Explain how missing data were addressed | 7-8 |
|  |  | (*d*) If applicable, describe analytical methods taking account of sampling strategy | 7-8 |
|  |  | (*e*) Describe any sensitivity analyses | 7-8 |
| **Results** |  |  |  |
| Participants | 13* | (a) Report numbers of individuals at each stage of study—eg numbers potentially eligible, examined for eligibility, confirmed eligible, included in the study,  completing follow-up, and analysed | 9-11 |
|  |  | (b) Give reasons for non-participation at each stage | NA |
|  |  | (c) Consider use of a flow diagram | - |
| Descriptive data | 14* | (a) Give characteristics of study participants (eg demographic, clinical, social) and  information on exposures and potential confounders | 9 |
|  |  | (b) Indicate number of participants with missing data for each variable of interest | NA |
| Outcome data | 15* | Report numbers of outcome events or summary measures | NA |
| Main results | 16 | (*a*) Give unadjusted estimates and, if applicable, confounder-adjusted estimates and their precision (eg, 95% confidence interval). Make clear which confounders were  adjusted for and why they were included | 9-11 |
|  |  | (*b*) Report category boundaries when continuous variables were categorized | NA |
|  |  | (*c*) If relevant, consider translating estimates of relative risk into absolute risk for a  meaningful time period | NA |
| Other analyses | 17 | Report other analyses done—eg analyses of subgroups and interactions, and  sensitivity analyses | 9-11 |
| **Discussion** |  |  |  |
| Key results | 18 | Summarise key results with reference to study objectives | 12-13 |
| Limitations | 19 | Discuss limitations of the study, taking into account sources of potential bias or  imprecision. Discuss both direction and magnitude of any potential bias | 14 |
| Interpretation | 20 | Give a cautious overall interpretation of results considering objectives, limitations,  multiplicity of analyses, results from similar studies, and other relevant evidence | 12-14 |
| Generalisability | 21 | Discuss the generalisability (external validity) of the study results | 14 |
| **Other information** |  |  |  |
| Funding | 22 | Give the source of funding and the role of the funders for the present study and, if  applicable, for the original study on which the present article is based |  |

*Give information separately for exposed and unexposed groups.

**Note:** An Explanation and Elaboration article discusses each checklist item and gives methodological background and published examples of transparent reporting. The STROBE checklist is best used in conjunction with this article (freely available on the Web sites of PLoS Medicine at [http://www.plosmedicine.org/,](http://www.plosmedicine.org/) Annals of Internal Medicine at [http://www.annals.org/,](http://www.annals.org/) and Epidemiology at [http://www.epidem.com/).](http://www.epidem.com/)) Information on the STROBE Initiative is available at [www.strobe-statement.org.](http://www.strobe-statement.org/)

Table S1. Complete results for mixed models including main effects of condition, biomarker, and interactions for both vergence and pupil features.

| **Feature** | **Biomarker** | **Effect** | **β** | **SE** | **p** |
| --- | --- | --- | --- | --- | --- |
| **Vergence** | | | | | |
| Slope Initial | CSF Aβ42 | Main effect: Condition | 0.0245 | 0.0396 | 0.536 |
|  |  | Main effect: Biomarker | -0.0199 | 0.0287 | 0.488 |
|  |  | Interaction | 0.0443 | 0.0398 | 0.265 |
| Slope Initial | p-Tau | Main effect: Condition | 0.0243 | 0.0396 | 0.539 |
|  |  | Main effect: Biomarker | -0.0575 | 0.0257 | 0.025 |
|  |  | Interaction | -0.0364 | 0.0388 | 0.349 |
| Slope Initial | Aβ42/p-Tau ratio | Main effect: Condition | 0.0244 | 0.0396 | 0.537 |
|  |  | Main effect: Biomarker | 0.0114 | 0.0277 | 0.680 |
|  |  | Interaction | 0.0009 | 0.0395 | 0.981 |
| Slope Global | CSF Aβ42 | Main effect: Condition | 0.0318 | 0.0391 | 0.416 |
|  |  | Main effect: Biomarker | -0.0307 | 0.0433 | 0.479 |
|  |  | Interaction | 0.0198 | 0.0393 | 0.615 |
| Slope Global | p-Tau | Main effect: Condition | 0.0319 | 0.0391 | 0.414 |
|  |  | Main effect: Biomarker | 0.0049 | 0.0426 | 0.908 |
|  |  | Interaction | -0.0492 | 0.0384 | 0.199 |
| Slope Global | Aβ42/p-Tau ratio | Main effect: Condition | 0.0319 | 0.0391 | 0.414 |
|  |  | Main effect: Biomarker | -0.0288 | 0.0412 | 0.484 |
|  |  | Interaction | -0.0251 | 0.0391 | 0.521 |
| Slope Late | CSF Aβ42 | Main effect: Condition | -0.0195 | 0.0398 | 0.623 |
|  |  | Main effect: Biomarker | 0.0232 | 0.0232 | 0.316 |
|  |  | Interaction | -0.0077 | 0.0400 | 0.847 |
| Slope Late | p-Tau | Main effect: Condition | -0.0195 | 0.0398 | 0.623 |
|  |  | Main effect: Biomarker | 0.0174 | 0.0231 | 0.453 |
|  |  | Interaction | 0.0101 | 0.0390 | 0.795 |
| Slope Late | Aβ42/p-Tau ratio | Main effect: Condition | -0.0197 | 0.0398 | 0.621 |
|  |  | Main effect: Biomarker | -0.0014 | 0.0229 | 0.952 |
|  |  | Interaction | 0.0240 | 0.0397 | 0.547 |
| AUC | CSF Aβ42 | Main effect: Condition | 0.0838 | 0.0390 | 0.031 |
|  |  | Main effect: Biomarker | -0.0303 | 0.0439 | 0.490 |
|  |  | Interaction | 0.0153 | 0.0392 | 0.696 |
| AUC | p-Tau | Main effect: Condition | 0.0840 | 0.0390 | 0.031 |
|  |  | Main effect: Biomarker | -0.0348 | 0.0424 | 0.412 |
|  |  | Interaction | -0.0426 | 0.0382 | 0.265 |
| AUC | Aβ42/p-Tau ratio | Main effect: Condition | 0.0840 | 0.0390 | 0.031 |
|  |  | Main effect: Biomarker | -0.0097 | 0.0422 | 0.818 |
|  |  | Interaction | -0.0371 | 0.0389 | 0.341 |
| Time to Peak | CSF Aβ42 | Main effect: Condition | -0.0004 | 0.0388 | 0.993 |
|  |  | Main effect: Biomarker | -0.0534 | 0.0470 | 0.257 |
|  |  | Interaction | 0.1084 | 0.0390 | 0.005 |
| Time to Peak | p-Tau | Main effect: Condition | -0.0001 | 0.0388 | 0.998 |
|  |  | Main effect: Biomarker | 0.0392 | 0.0461 | 0.395 |
|  |  | Interaction | -0.0872 | 0.0381 | 0.022 |
| Time to Peak | Aβ42/p-Tau ratio | Main effect: Condition | -0.0005 | 0.0389 | 0.989 |
|  |  | Main effect: Biomarker | -0.0408 | 0.0449 | 0.364 |
|  |  | Interaction | 0.0606 | 0.0388 | 0.119 |
| Peak Power | CSF Aβ42 | Main effect: Condition | 0.0112 | 0.0354 | 0.752 |
|  |  | Main effect: Biomarker | -0.0254 | 0.0827 | 0.759 |
|  |  | Interaction | 0.0402 | 0.0355 | 0.258 |
| Peak Power | p-Tau | Main effect: Condition | 0.0115 | 0.0353 | 0.746 |
|  |  | Main effect: Biomarker | -0.0170 | 0.0803 | 0.832 |
|  |  | Interaction | -0.0859 | 0.0347 | 0.013 |
| Peak Power | Aβ42/p-Tau ratio | Main effect: Condition | 0.0111 | 0.0354 | 0.753 |
|  |  | Main effect: Biomarker | 0.0078 | 0.0786 | 0.921 |
|  |  | Interaction | 0.0103 | 0.0353 | 0.772 |
| **Pupil** | | | | | |
| Slope Initial | CSF Aβ42 | Main effect: Condition | 0.0567 | 0.0396 | 0.152 |
|  |  | Main effect: Biomarker | -0.0197 | 0.0265 | 0.458 |
|  |  | Interaction | 0.0007 | 0.0398 | 0.986 |
| Slope Initial | p-Tau | Main effect: Condition | 0.0568 | 0.0396 | 0.152 |
|  |  | Main effect: Biomarker | 0.0321 | 0.0263 | 0.223 |
|  |  | Interaction | -0.0256 | 0.0389 | 0.509 |
| Slope Initial | Aβ42/p-Tau ratio | Main effect: Condition | 0.0567 | 0.0396 | 0.152 |
|  |  | Main effect: Biomarker | -0.0392 | 0.0252 | 0.120 |
|  |  | Interaction | 0.0067 | 0.0396 | 0.865 |
| Slope Global | CSF Aβ42 | Main effect: Condition | 0.7245 | 0.0358 | <0.001 |
|  |  | Main effect: Biomarker | 0.0741 | 0.0593 | 0.211 |
|  |  | Interaction | -0.0580 | 0.0360 | 0.108 |
| Slope Global | p-Tau | Main effect: Condition | 0.7248 | 0.0358 | <0.001 |
|  |  | Main effect: Biomarker | -0.1307 | 0.0528 | 0.013 |
|  |  | Interaction | -0.0724 | 0.0352 | 0.040 |
| Slope Global | Aβ42/p-Tau ratio | Main effect: Condition | 0.7245 | 0.0359 | <0.001 |
|  |  | Main effect: Biomarker | 0.0942 | 0.0551 | 0.087 |
|  |  | Interaction | -0.0197 | 0.0358 | 0.582 |
| Slope Late | CSF Aβ42 | Main effect: Condition | -0.0448 | 0.0396 | 0.259 |
|  |  | Main effect: Biomarker | -0.0171 | 0.0265 | 0.517 |
|  |  | Interaction | -0.0764 | 0.0398 | 0.055 |
| Slope Late | p-Tau | Main effect: Condition | -0.0448 | 0.0396 | 0.258 |
|  |  | Main effect: Biomarker | 0.0301 | 0.0263 | 0.253 |
|  |  | Interaction | 0.0197 | 0.0389 | 0.613 |
| Slope Late | Aβ42/p-Tau ratio | Main effect: Condition | -0.0443 | 0.0396 | 0.264 |
|  |  | Main effect: Biomarker | -0.0151 | 0.0256 | 0.555 |
|  |  | Interaction | -0.0919 | 0.0396 | 0.020 |
| AUC | CSF Aβ42 | Main effect: Condition | 0.5606 | 0.0371 | <0.001 |
|  |  | Main effect: Biomarker | 0.0409 | 0.0548 | 0.455 |
|  |  | Interaction | -0.0250 | 0.0373 | 0.502 |
| AUC | p-Tau | Main effect: Condition | 0.5608 | 0.0370 | <0.001 |
|  |  | Main effect: Biomarker | -0.0682 | 0.0519 | 0.188 |
|  |  | Interaction | -0.0735 | 0.0364 | 0.043 |
| AUC | Aβ42/p-Tau ratio | Main effect: Condition | 0.5605 | 0.0371 | <0.001 |
|  |  | Main effect: Biomarker | 0.0243 | 0.0524 | 0.643 |
|  |  | Interaction | 0.0134 | 0.0370 | 0.718 |
| Time to Peak | CSF Aβ42 | Main effect: Condition | 0.5870 | 0.0371 | <0.001 |
|  |  | Main effect: Biomarker | 0.0609 | 0.0513 | 0.235 |
|  |  | Interaction | 0.0138 | 0.0373 | 0.711 |
| Time to Peak | p-Tau | Main effect: Condition | 0.5874 | 0.0370 | <0.001 |
|  |  | Main effect: Biomarker | -0.0427 | 0.0499 | 0.391 |
|  |  | Interaction | -0.1289 | 0.0364 | <0.001 |
| Time to Peak | Aβ42/p-Tau ratio | Main effect: Condition | 0.5867 | 0.0371 | <0.001 |
|  |  | Main effect: Biomarker | 0.0537 | 0.0486 | 0.269 |
|  |  | Interaction | 0.0729 | 0.0370 | 0.049 |
| Peak Power | CSF Aβ42 | Main effect: Condition | 0.5918 | 0.0364 | <0.001 |
|  |  | Main effect: Biomarker | -0.0085 | 0.0596 | 0.887 |
|  |  | Interaction | 0.0434 | 0.0366 | 0.236 |
| Peak Power | p-Tau | Main effect: Condition | 0.5925 | 0.0362 | <0.001 |
|  |  | Main effect: Biomarker | 0.0214 | 0.0580 | 0.712 |
|  |  | Interaction | -0.2392 | 0.0356 | <0.001 |
| Peak Power | Aβ42/p-Tau ratio | Main effect: Condition | 0.5913 | 0.0364 | <0.001 |
|  |  | Main effect: Biomarker | -0.0131 | 0.0567 | 0.817 |
|  |  | Interaction | 0.1150 | 0.0364 | 0.002 |

All models included sex, MMSE and age as covariates. β represents standardized regression coefficients. SE = standard error. Condition effect represents the difference between Target and Distractor trials. Biomarker effect represents the main effect of the respective CSF biomarker (Aβ42, pTau, or Aβ42/pTau ratio). Interaction represents the Condition × Biomarker interaction term.

Table S2. Complete results for Linear models.

| **Feature** | **Biomarker** | **β** | **SE** | **p-value** | **R²** |
| --- | --- | --- | --- | --- | --- |
| **Vergence** | | | | | |
| Slope Initial | pTau | -0.784 | 0.315 | 0.015 | 0.20 |
|  | Aβ42 | -0.253 | 0.333 | 0.449 | 0.09 |
|  | Aβ42/pTau | 0.169 | 0.337 | 0.619 | 0.06 |
| Slope Global | pTau | -0.069 | 0.271 | 0.801 | 0.10 |
|  | Aβ42 | -0.177 | 0.273 | 0.521 | 0.09 |
|  | Aβ42/pTau | -0.174 | 0.276 | 0.532 | 0.07 |
| Slope Late | pTau | -0.044 | 0.206 | 0.831 | 0.11 |
|  | Aβ42 | 0.168 | 0.208 | 0.421 | 0.09 |
|  | Aβ42/pTau | 0.077 | 0.211 | 0.716 | 0.07 |
| Auc | pTau | -0.366 | 0.311 | 0.243 | 0.12 |
|  | Aβ42 | -0.193 | 0.317 | 0.544 | 0.09 |
|  | Aβ42/pTau | -0.031 | 0.320 | 0.924 | 0.07 |
| Time To Peak | pTau | 0.072 | 0.208 | 0.732 | 0.10 |
|  | Aβ42 | -0.146 | 0.209 | 0.487 | 0.09 |
|  | Aβ42/pTau | -0.104 | 0.212 | 0.625 | 0.06 |
| Peak Power | pTau | -0.105 | 0.220 | 0.635 | 0.11 |
|  | Aβ42 | -0.054 | 0.223 | 0.811 | 0.08 |
|  | Aβ42/pTau | 0.051 | 0.226 | 0.821 | 0.06 |
| **Pupil** | | | | | |
| Slope Initial | pTau | 0.077 | 0.166 | 0.645 | 0.10 |
|  | Aβ42 | -0.032 | 0.168 | 0.850 | 0.09 |
|  | Aβ42/pTau | -0.109 | 0.169 | 0.521 | 0.07 |
| Slope Global | pTau | -0.610 | 0.277 | 0.016 | 0.21 |
|  | Aβ42 | 0.358 | 0.291 | 0.224 | 0.10 |
|  | Aβ42/pTau | 0.458 | 0.292 | 0.121 | 0.10 |
| Slope Late | pTau | 0.318 | 0.351 | 0.368 | 0.14 |
|  | Aβ42 | -0.302 | 0.358 | 0.402 | 0.10 |
|  | Aβ42/pTau | -0.271 | 0.359 | 0.454 | 0.10 |
| Auc | pTau | -0.294 | 0.244 | 0.233 | 0.15 |
|  | Aβ42 | 0.213 | 0.252 | 0.401 | 0.09 |
|  | Aβ42/pTau | 0.135 | 0.255 | 0.598 | 0.07 |
| Time To Peak | pTau | -0.197 | 0.276 | 0.478 | 0.15 |
|  | Aβ42 | 0.325 | 0.282 | 0.254 | 0.10 |
|  | Aβ42/pTau | 0.272 | 0.284 | 0.342 | 0.10 |
| Peak Power | pTau | 0.135 | 0.367 | 0.715 | 0.13 |
|  | Aβ42 | -0.024 | 0.377 | 0.950 | 0.08 |
|  | Aβ42/pTau | -0.089 | 0.380 | 0.815 | 0.07 |

All models included sex, MMSE and age as covariates. β represents standardized regression coefficients. SE = standard error.
